# Prioritisation of Structural Variant Calls in Cancer Genomes

**DOI:** 10.1101/084640

**Authors:** Miika J Ahdesmäki, Brad Chapman, Pablo E Cingolani, Oliver Hofmann, Aleksandr Sidoruk, Zhongwu Lai, Gennadii Zakharov, Mikhail Rodichenko, Mikhail Alperovich, David Jenkins, T. Hedley Carr, Daniel Stetson, Brian Dougherty, J. Carl Barrett, Justin J Johnson

**Affiliations:** AstraZeneca Oncology IMed; Harvard T.H. Chan School of Public Health; Kew Inc.; University of Melbourne Centre for Cancer Research; EPAM Systems Inc.; Boston University

## Abstract

Sensitivity of short read DNA-sequencing for gene fusion detection is improving, but is hampered by the significant amount of noise composed of uninteresting or false positive hits in the data. In this paper we describe a tiered prioritisation approach to extract high impact gene fusion events. Using cell line and patient DNA sequence data we improve the annotation and interpretation of structural variant calls to best highlight likely cancer driving fusions. We also considerably improve on the automated visualisation of the high impact structural variants to highlight the effects of the variants on the resulting transcripts. The resulting framework greatly improves on readily detecting clinically actionable structural variants.

## Introduction

Structural variants (SVs) such as inversions, tandem duplications, large deletions and more complex chromosomal rearrangements are implicated as driver events in multiple cancers [Latysheva]. The mechanisms for oncogenic driver generation include activating fusions combining the coding frames (quite often in the intronic regions) of two genes, as well as truncating mutations in tumor suppressor genes or whole exon losses. Some well understood examples include TMPRSS2-ERG in prostate cancer [Tomlins], FGFR1/3-TACC1/3 in bladder and other cancers [Weinstein], EGFRv3 deletion in glioblastoma and other tumours [Sugawa] and EML4-ALK in lung cancer [Soda]. Clinical detection of SVs in Mendelian diseases has been considered by others, see [Noll].

The accurate calling of these complex, structural variants in short read DNA sequencing data is complicated by noise, manifested as false positives and lack of specificity. In many cases, the number of whole genome SV calls, including complex breakends, can be in the tens of thousands. While long read sequencing is likely to improve the calling of structural variants especially in germline DNA, tumour DNA in FFPE and circulating tumour DNA samples is inherently limited to short DNA fragment size. Utilising the currently available large amount of short read sequencing data to the full is therefore well motivated. It is also imperative to promptly pinpoint any clinically important structural variants when present in data.

In this paper we propose a tiered prioritisation approach to extract structural variants most likely to contribute to cancer proliferation and enable validation and follow up for a subset of high priority events. The prioritisation is based on greatly improved structural variant annotation in the variant annotation tool SnpEff [Cingolani]. Similar prioritisation work has been published in the domain of small variants, see for example [Carr, Muenz]. The focus here shall be on DNA data only, as fusion calling and prioritisation is well established in RNA-seq data, see e.g. https://github.com/ndaniel/fusioncatcher [Nicorici].

The important aspect of easy and automated visualisation of the effects of structural variants on genes and coding exons is often overlooked with focus on structural variant calling algorithm performance. We thus further implement interactive structural variant visualisations in the New Genome Browser. We show the full utility of the improved prioritisation and visualisation approaches in samples with structural variants leading to oncogenic gene fusions.

## Methods

Most short read SV calling pipelines start with alignment of the DNA data to the human reference using an aligner like bwa-mem [Li]. This is followed by integrating evidence from split and discordant reads, and potentially coverage [Alkan], to make structural variant calls for deletions (DEL), tandem duplications (DUP), inversions (INV) and other more complex variants (BND). An example of these events is visualised in Figure 1 in [Tattini].

**Figure 1.**
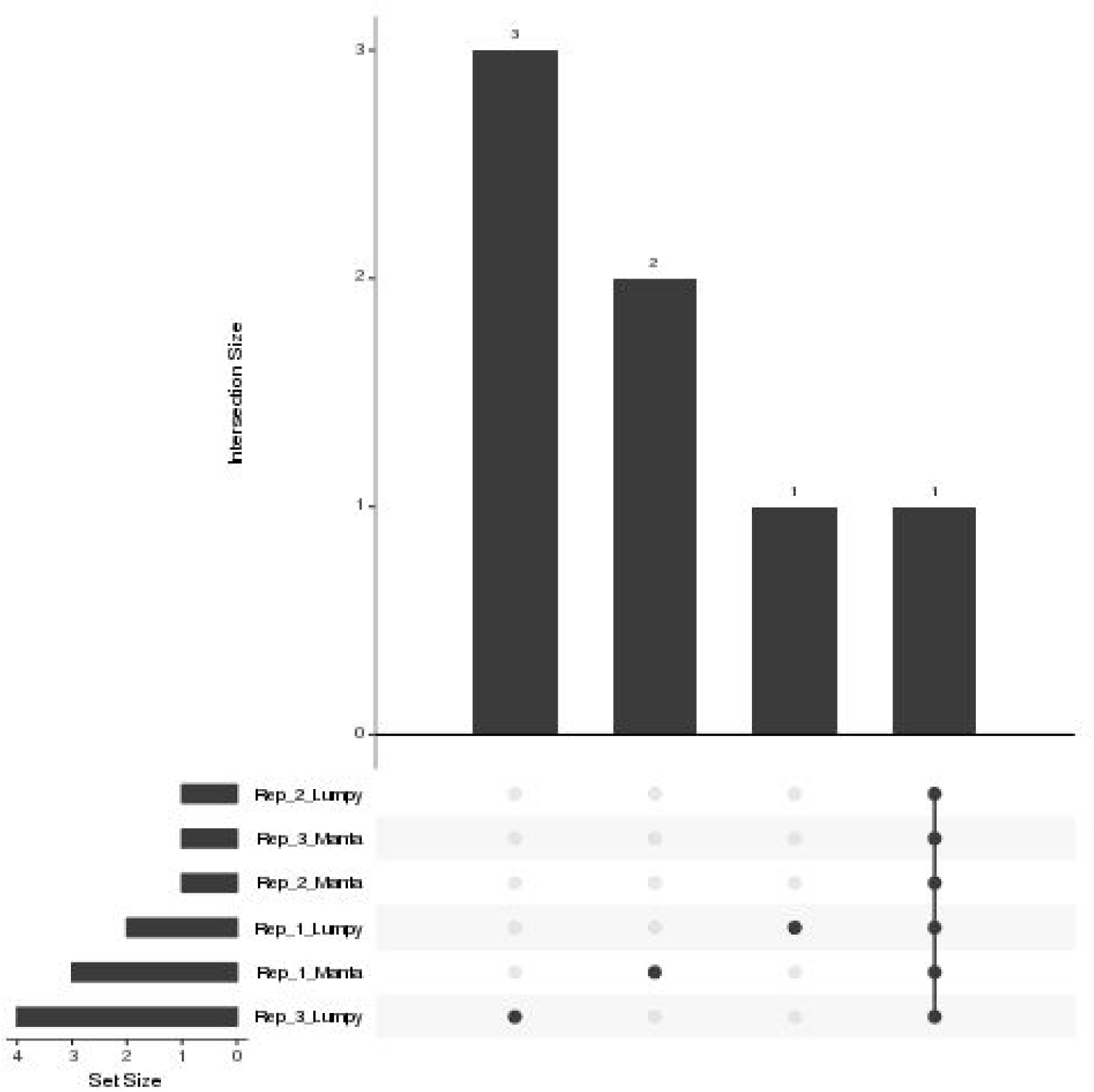
HDC134P prioritised SV call concordance. One event (EML4-ALK) is detected by all the callers and several other private events (false positives) are detected in some of the replicates.

**Figure 2.**
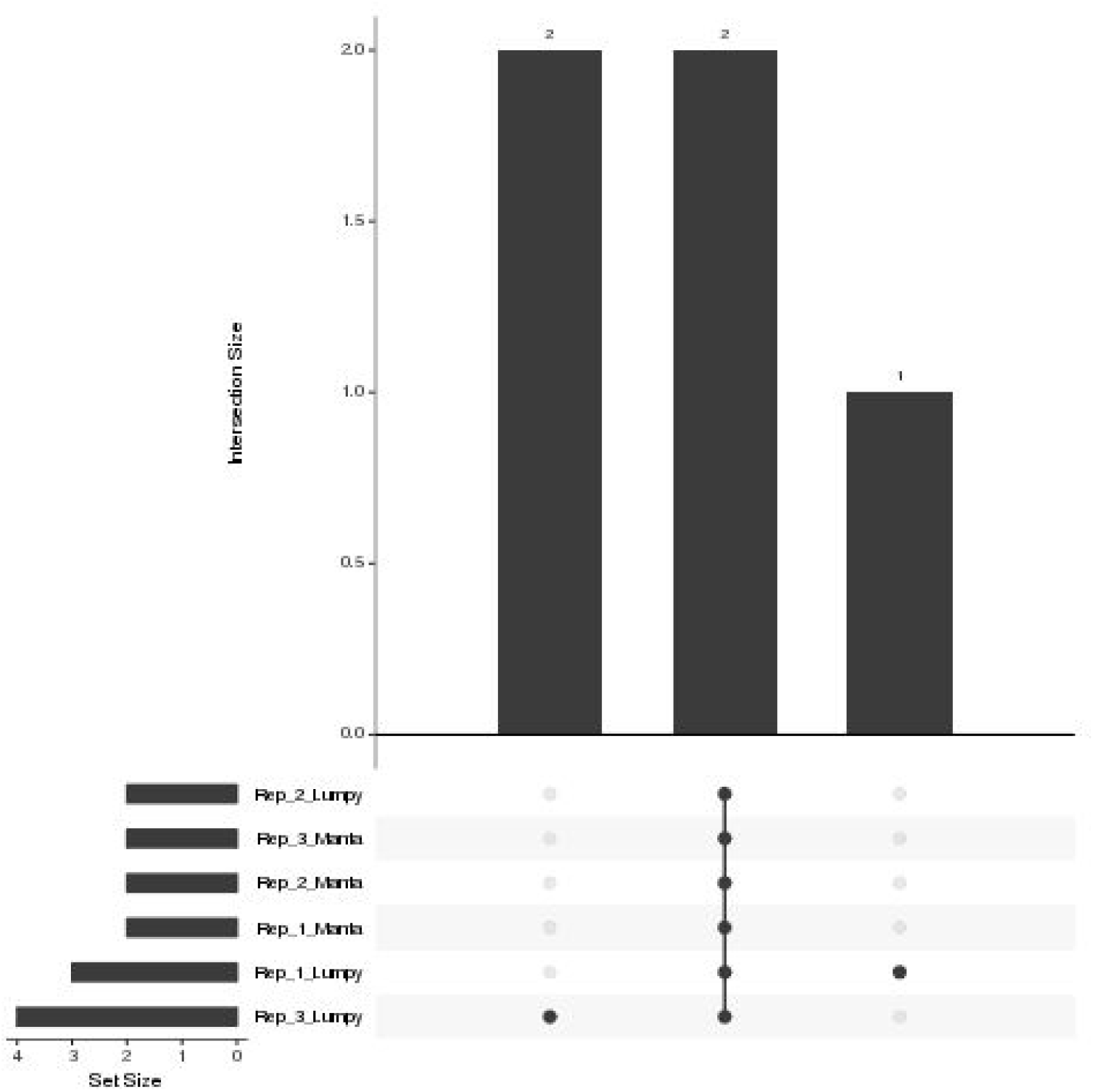
HDC140P prioritised SV call concordance. Two events (CCDC6-RET and RET-chr13) are detected by all the callers and 3 private events (false positives) are detected in some of the replicates.

**Figure 3.**
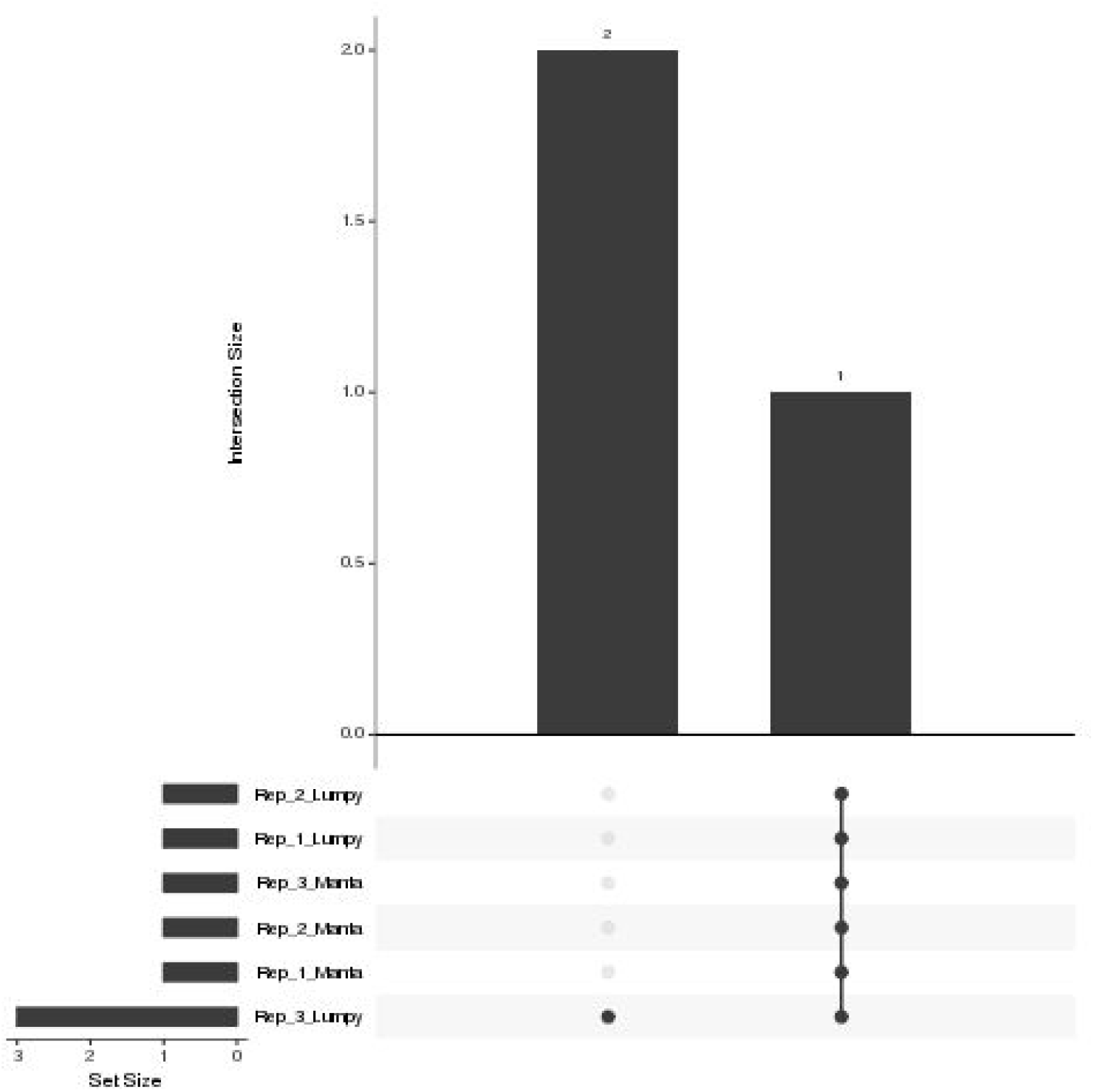
HDC141P prioritised SV call concordance. One event (SLC34A2-ROS1) is detected by all the callers and two other private events (false positives) are detected in some of the replicates.

We utilised two freely available SV callers that integrate evidence from split and discordant reads, Manta [Chen] and Lumpy [Layer]. Both benchmark well in synthetic somatic data sets (see ICGC-TCGA DREAM Mutation Calling challenge leaderboards https://www.synapse.org/#!Synapse:syn312572/wiki/247695) as well as germline reference standards (Genome in a Bottle NA12878). Any structural variant caller (such as BRASS, https://github.com/cancerit/BRASS) producing vcf files compliant with the vcf specification (https://samtools.github.io/hts-specs/VCFv4.3.pdf) and compatible with SnpEff could equally well be used with our proposed methodology, provided they also quantify the numbers of discordant and split reads supporting at least the alternative allele.

We built a three tier prioritisation system (https://github.com/AstraZeneca-NGS/simple_sv_annotation) using fusion and exon loss annotations. Given a list of genes of interest (GOI):

- Gene fusion

- Fusion affecting two genes

§ on list of known pairs from FusionCatcher [Nicorici] (1)
§ not on list of known pairs

- one or two genes on gene list (2)
- neither gene on gene list (3)
- Fusion affecting one gene

§ on gene list (2)
§ not on gene list (3)
- Whole exon loss

- on prioritisation gene list (2)
- not on gene list (3)
- Upstream or downstream of gene list genes (3)
- Other variant (REJECT)
- Missing ANN or SVTYPE in variant call file (REJECT)

For the GOI, we have provided a list of 300+ genes commonly associated with cancer, including genes involved in the MAPK and PI3K pathways (including receptor tyrosine kinase genes), DNA damage response, immuno-oncology and others. Alternatively, the user can provide their own gene lists in the implementation. The proposed prioritisation approach can be applied to variants from both paired (tumour/normal) and tumour only data, depending only on the structural variant callers’ capabilities to handle paired samples. We confirmed the approach using TCGA data with known gene fusions.

To practically facilitate the prioritisation we improved annotations in SnpEff 4.3 to ease interpretation of fusion events, adding the Sequence Ontology [Eilbeck] annotation type *gene_fusion* for events where the open reading frames are in the same direction. Further, *bidirectional_gene_fusion* was introduced for where the frames of the putatively fused genes are opposing and therefore unlikely to be functional and *frameshift_variant* when the coding of the resulting fusion is out of frame, thus likely resulting in a truncated protein. The last two types are very important and interesting for loss of function of e.g. tumour suppressors. Other annotation improvements in SnpEff 4.3 include: *chromosome_number_variation, duplication and inversion*, which refer to large chromosomal deletions, duplications and inversions respectively (involving a whole exon, transcript, gene or even larger genomic regions), *exon_loss_variant* (whole or significant part of the exon was deleted) and *feature_ablation* (whole gene deleted).

## Results

We estimated the ability of our prioritisation approach to retain known mutations while reducing false positive events using samples with known structural variants (in a background of less well characterised SVs). Synthetic datasets from the ICGC-TCGA DREAM Mutation Calling challenge have also known artificial structural variants spiked in. These artificial SVs are useful, but do not however necessarily represent a realistic quantity of somatic SVs since they are not generated from a biological model, so we focused on known events in real sequenced samples.

We collected sequencing data for seven samples with known SVs from cell lines, a patient derived xenograft and a clinical sample (Table 1). Although whole genome sequencing data or targeted capture including introns is preferred, any hybrid capture data can yield meaningful results if the breakpoints are close to captured regions or there are off target reads.

**Table 1.** Collection of structural variants leading to oncogenic fusions in different sample types. All events are ranked into the highest category (1). * Data for the patient derived xenograft (PDX) model could not be shared.

| Sample | Panel or WGS | Manta call | Lumpy call | Fusion |
| --- | --- | --- | --- | --- |
| HDC134P replicate 1 | Panel with intronic probes | INV | INV | EML4-ALK |
| HDC134P replicate 2 | Panel with intronic probes | INV | INV | EML4-ALK |
| HDC134P replicate 3 | Panel with intronic probes | INV | BND | EML4-ALK |
| HDC140P replicate 1 | Panel with intronic probes | INV | BND | CCDC6-RET |
| HDC140P replicate 2 | Panel with intronic probes | INV | BND | CCDC6-RET |
| HDC140P replicate 3 | Panel with intronic probes | INV | BND | CCDC6-RET |
| HDC141P replicate 1 | Panel with intronic probes | BND | BND | ROS1-SLC34A2 |
| HDC141P replicate 2 | Panel with intronic probes | BND | BND | ROS1-SLC34A2 |
| HDC141P replicate 3 | Panel with intronic probes | BND | BND | ROS1-SLC34A2 |
| MCF7 | WGS | DUP | BND, DUP | RAD51C-ATXN7, ESR1-CCDC170 |
| RT4 | Panel with intronic probes | DUP | DUP | TACC3-FGFR3 |
| PDX model* | WES | DUP | DUP | TACC3-FGFR3 |
| Prostate cancer patient sample | FMI Panel with intronic probes | DEL | DEL | TMPRSS2-ERG |

Following bwa-mem alignment to hg38 Lumpy and Manta both call the breakpoints and event types for the structural variants in Table 1, with slightly different interpretations for some like CCDC6-RET. As part of the updates to SnpEff we ensured that all these different types of SV events (INV, DUP, DEL) affecting two genes were correctly annotated as gene fusions.

The total number of calls for the samples in Table 1 as well as the numbers of variants falling into the tiers are shown in Table 2 where it is evident that Lumpy produces more calls particularly when compared to Manta. We recommend Manta 1.0 or later for good sensitivity and fewer (likely) false positives compared to Lumpy.

**Table 2.** Raw SV call numbers for Manta and Lumpy are given in the DUP, DEL, INV and BND columns. The prioritised calls are shown in the last three columns. The Primary priority column corresponds to the number of detected fusions reported previously in the literature. All samples are from small hybrid capture panels except for the MCF7 sample, thus the relatively low numbers of calls per sample.

| Sample | Algorithm | DUP, DEL, INV | BND | Primary priority | Secondary priority | Tertiary priority |
| --- | --- | --- | --- | --- | --- | --- |
| HDC134P replicate 1 | Manta | 5 | 0 | 2 | 2 | 0 |
|  | Lumpy | 43 | 9 | 1 | 0 | 1 |
| HDC134P replicate 2 | Manta | 2 | 0 | 2 | 0 | 0 |
|  | Lumpy | 41 | 4 | 1 | 0 | 0 |
| HDC134P replicate 3 | Manta | 3 | 0 | 2 | 0 | 0 |
|  | Lumpy | 41 | 2 | 1 | 2 | 1 |
| HDC140P replicate 1 | Manta | 1 | 1 | 1 | 1 | 0 |
|  | Lumpy | 48 | 20 | 1 | 2 | 0 |
| HDC140P<br>replicate 2 | Manta | 1 | 1 | 1 | 1 | 0 |
|  | Lumpy | 30 | 12 | 1 | 1 | 0 |
| HDC140P<br>replicate 3 | Manta | 1 | 1 | 1 | 1 | 0 |
|  | Lumpy | 56 | 9 | 1 | 3 | 0 |
| HDC141P<br>replicate 1 | Manta | 0 | 2 | 2 | 0 | 0 |
|  | Lumpy | 24 | 7 | 1 | 0 | 0 |
| HDC141P<br>replicate 2 | Manta | 0 | 2 | 2 | 0 | 0 |
|  | Lumpy | 17 | 1 | 1 | 0 | 0 |
| HDC141P<br>replicate 3 | Manta | 0 | 2 | 2 | 0 | 0 |
|  | Lumpy | 25 | 6 | 1 | 1 | 1 |
| MCF7<br>(WGS) | Manta | 8239 | 1990 | 16 | 48 | 2814 |
|  | Lumpy | 4277 | 2750 | 21 | 38 | 1683 |
| RT4 | Manta | 169 | 158 | 1 | 20 | 135 |
|  | Lumpy | 1509 | 14659 | 1 | 302 | 10300 |
| PDX<br>model | Manta | 248 | 65 | 5 | 2 | 164 |
|  | Lumpy | 143 | 862 | 5 | 8 | 292 |
| Patient<br>sample | Manta | 30 | 51 | 1 | 6 | 37 |
|  | Lumpy | 1034 | 3621 | 1 | 13 | 177 |

In Table 2, the primary priority (known fusions) column lists the one true fusion known to be present in the HDC, RT4 cell lines and the patient sample. Manta reported two close but different breakpoints for the EML4-ALK fusion in the HDC134P replicates. The background of the PDX model is not fully characterised and therefore in the list of 5 fusions there may be false positives or SVs of unknown significance besides the FGFR3-TACC3 fusion. MCF7 has been previously characterised to at least contain the RAD51C-ATXN7 and ESR1-CCDC170 fusions [Hampton, Veeraraghavan]. The remaining >10 fusions may be false positives or of less importance. A large reduction in the amount of noise can be seen through the primary and secondary priority categories whereas the tertiary priority (upstream, downstream events in genes of interest and fusions in genes of uninterest) is a catch-all category that should receive less attention.

To visualise the prioritised SV calls in the three replicate samples (HDC134P, HDC140P, HDC141P) run in triplicates, we utilised the UpSet package [Lex] to show con- and discordance in the prioritised calls. The plots in Figure 1 through 3 show the concordance histograms for each of the triplicates. The known fusions are detected by both Lumpy and Manta in all triplicates. There is additionally one structural variant (RET fused with chromosome 13) in HDC140P detected by all the algorithms in all the replicates. All the rest of the calls are private to one caller and one replicate (noise) but the number is small. This shows that the true events are very confidently called by both algorithms but there is a varying amount of false positives with Lumpy producing slightly more.

To show that the proposed approach correctly identifies the true events also in data not part of the sample set in Table 1, we applied the prioritisation to the TCGA bladder cancer cohort [Weinstein]. The FGFR3-TACC3 fusion in the RT4 cell line and PDX model in Table 1 is found in several TCGA samples, e.g. TCGA-CF-A3MF, TCGA-CF-A3MG and TCGA-CF-A3MH. Typically the breakpoints of gene fusions in the intronic regions, however in two of the TCGA samples (TCGA-CF-A3MH, TCGA-CF-A3MF), the FGFR3 breakpoint is in the last exon. The events were correctly annotated and identified in these samples by our approach.

### Visualising gene fusions resulting from structural variants

Visualisation of structural variants to highlight the breakpoints and affected exons in a putative fusion transcript is an area of active development with no one tool currently being the industry standard. We first utilised Svviz [Spies], one of the earlier tools, to examine the validated fusion variants identified by prioritisation. The FGFR3-TACC3 tandem duplication (RT4 cell line) is shown in Figure 4; TACC3 is not captured by the panel used and therefore no reads in support of the reference allele for TACC3 are shown. Svviz reassembles the reads around the putative breakpoints in its analysis and requires an amount of manual intervention.

**Figure 4.**
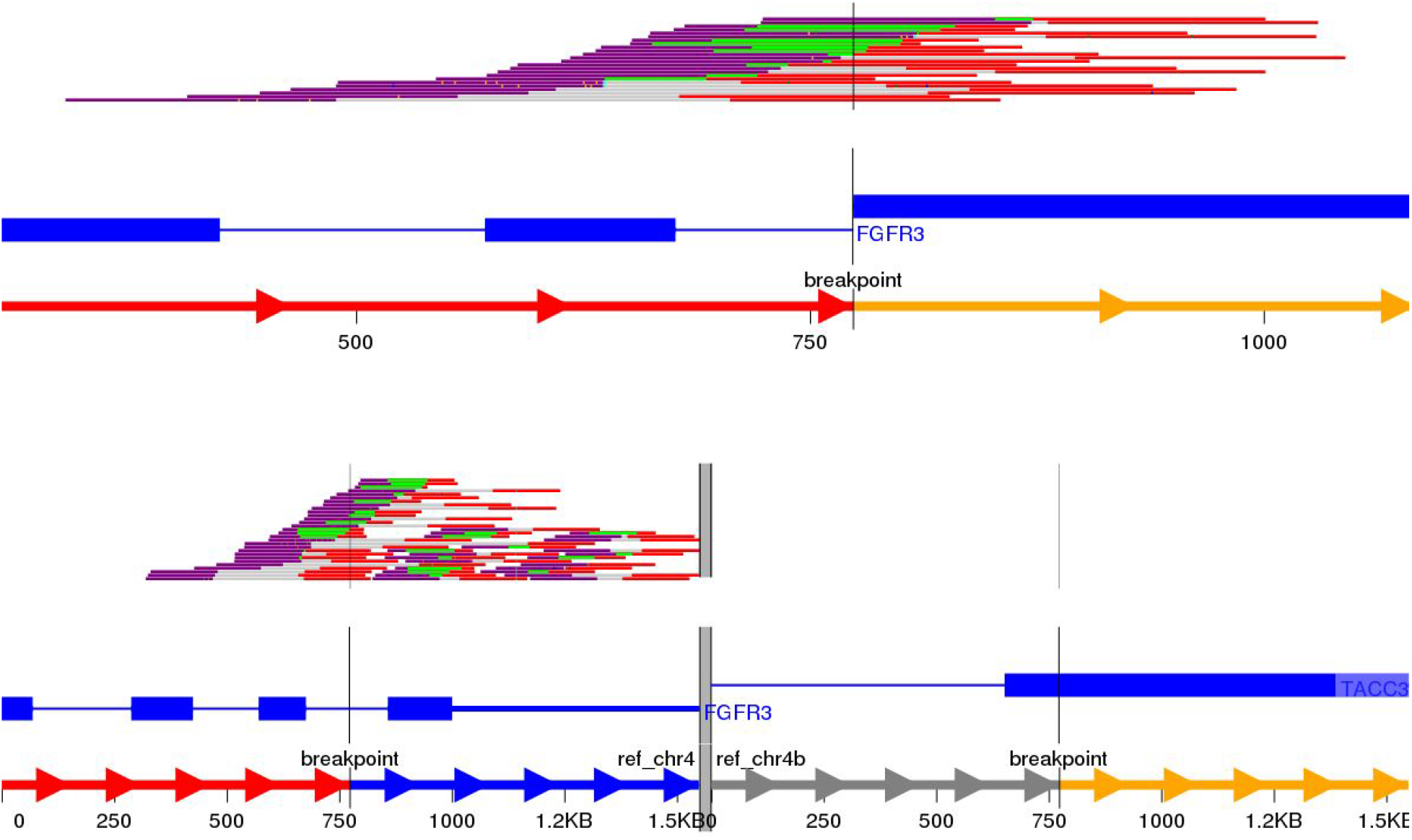
Svviz output for the FGFR3-TACC3 fusion (tandem duplication) in the RT4 cell line. The top row illustrates how the last intron of FGFR3 is fused to an exon of TACC3. The bottom row shows the read evidence for the reference alleles.

We next decided to implement a variant call based gene fusion visualisation scheme in the open source New Genome Browser (NGB, https://github.com/epam/NGB). NGB takes the variant breakpoints and uses Ensembl and UniProt based annotation to visualise the fusion product in both reference as well as the actual sequence context. The resulting plots highlight the fused exons of the affected genes.

The NGB visualisation of the FGFR3-TACC3 fusion is shown in Figure 5. Unlike the Svviz plot (figure 4), the visualisation is fully interactive html5 in the browser. Red highlighting is used to show the breakpoints relative to the coding regions in the alternative allele view and the red line shows the fusion points in the reference allele view.

**Figure 5.**
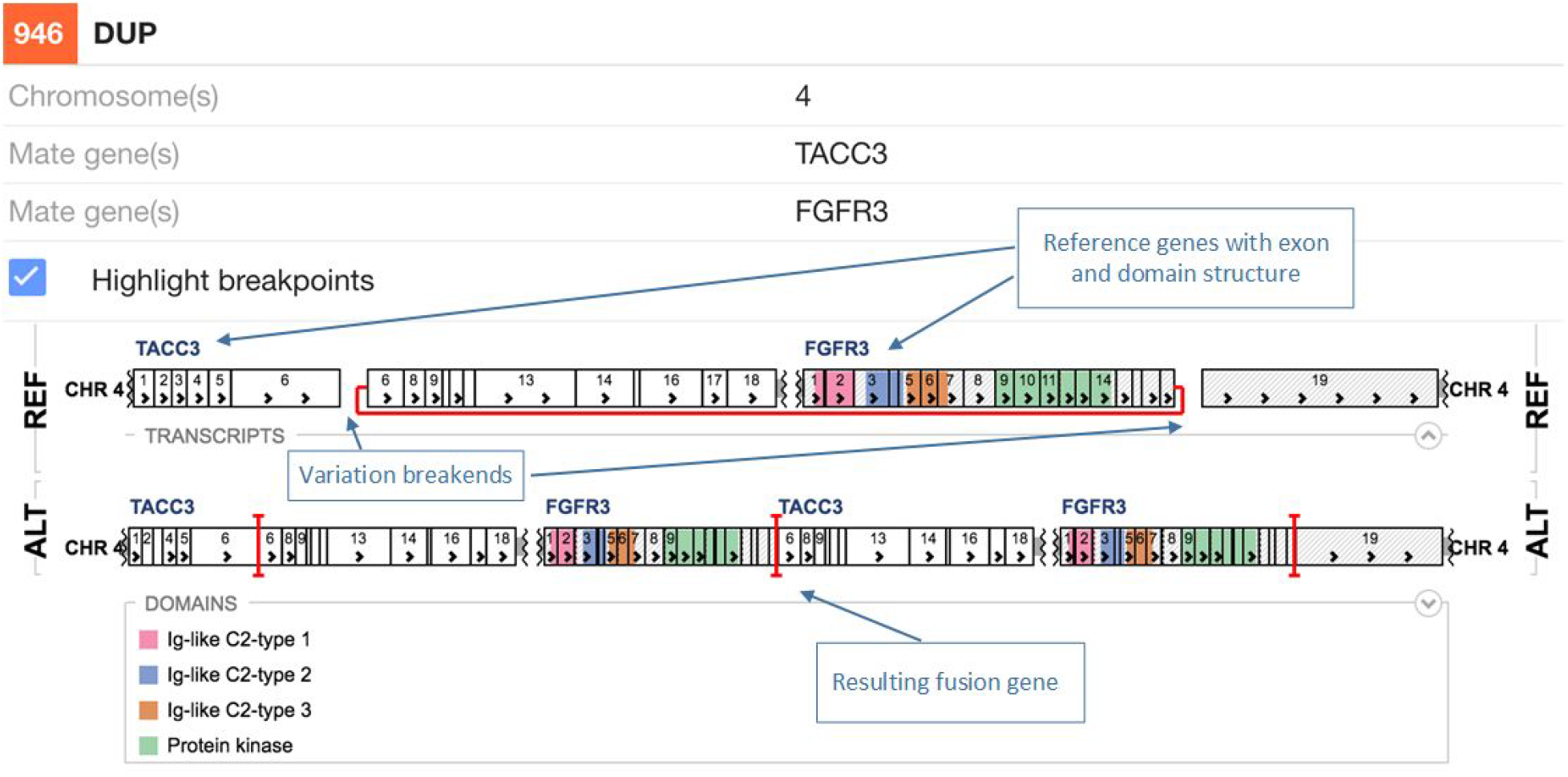
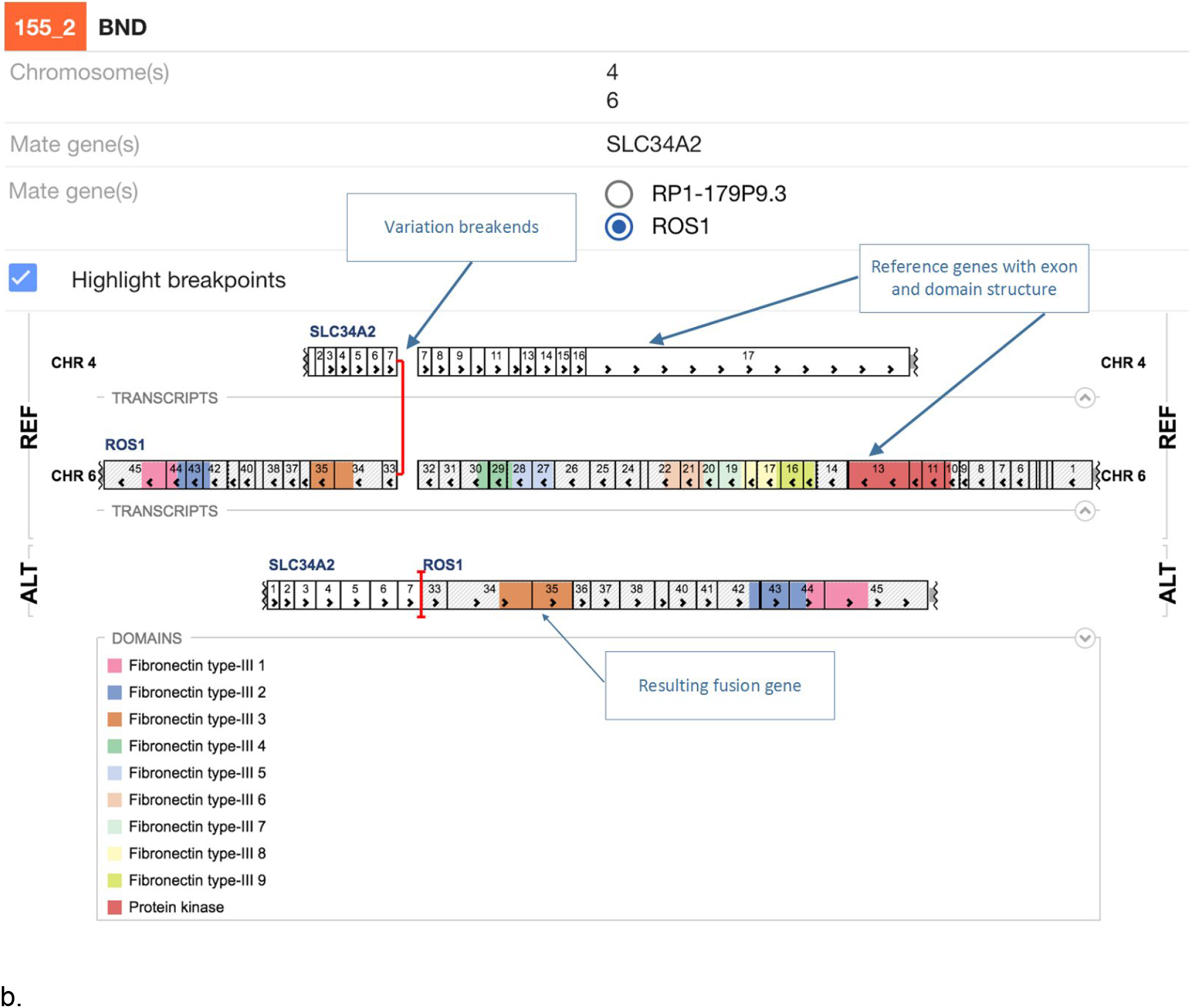
FGFR3-TACC3 tandem duplication fusion exon level visualisation in the New Genome Browser. Protein domains and exons affected by the structural variant are highlighted in colours.

As NGB is a full feature genome browser, viewing both the read evidence as well as the fusion effects is simple. In Figure 6a we shown an interchromosomal translocation resulting in a fusion between ROS1 and SLC34A2. If multiple genes are overlapping the breakpoints NGB allows choosing the most relevant gene for the researcher. figure 6b shows the read level evidence side-by-side from the two breakpoints. Soft clipping of the reads around the breakpoint are shown by the coloured base tails of the reads
a.

**Figure 6:**
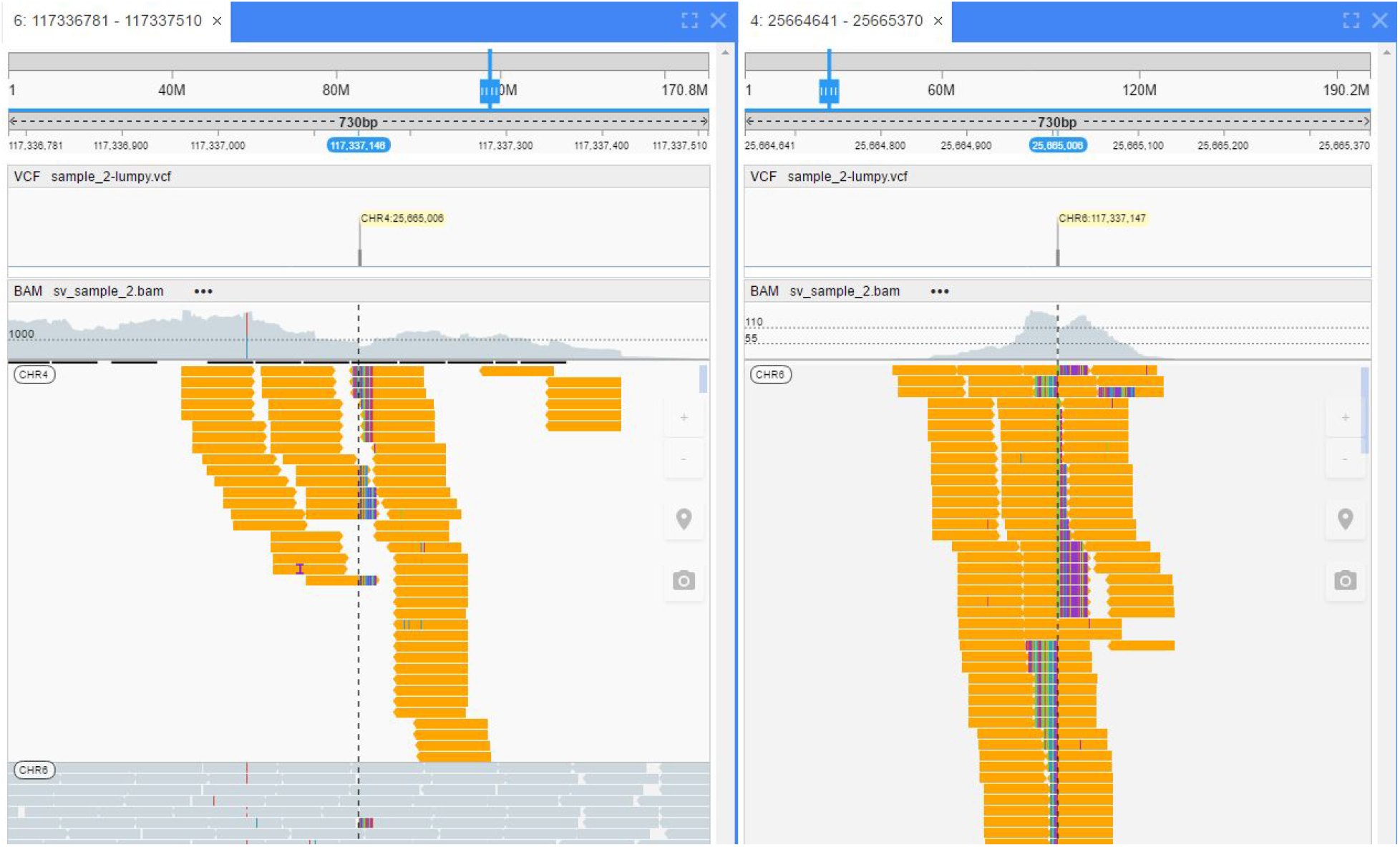
ROS1-SLC34A2 interchromosomal translocation fusion. a. The effect of the fusion. b. The read evidence for the event at both breakpoints.

Another example for EML4-ALK fusion the results from an inversion is shown if Figure 7.

**Figure 7.**
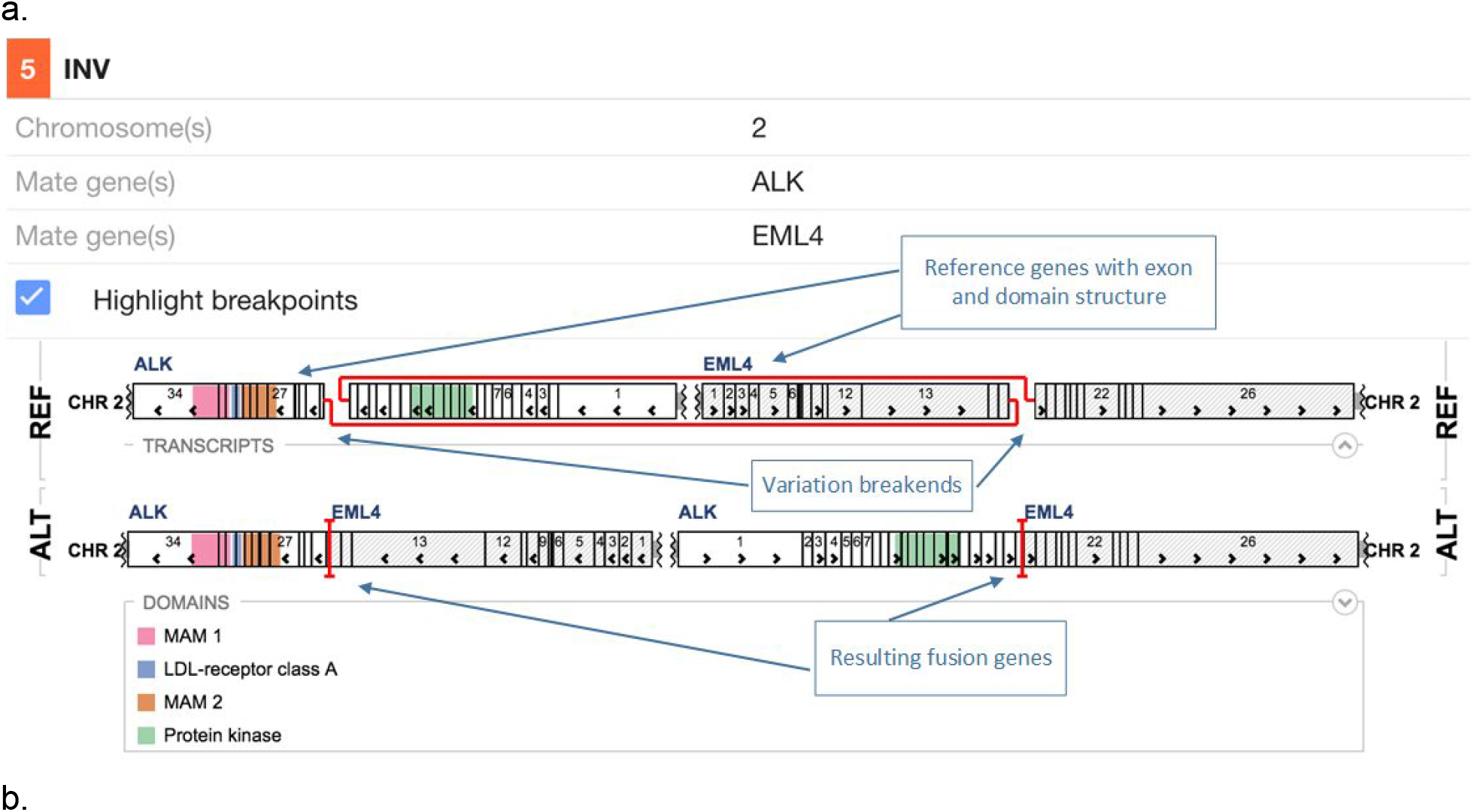
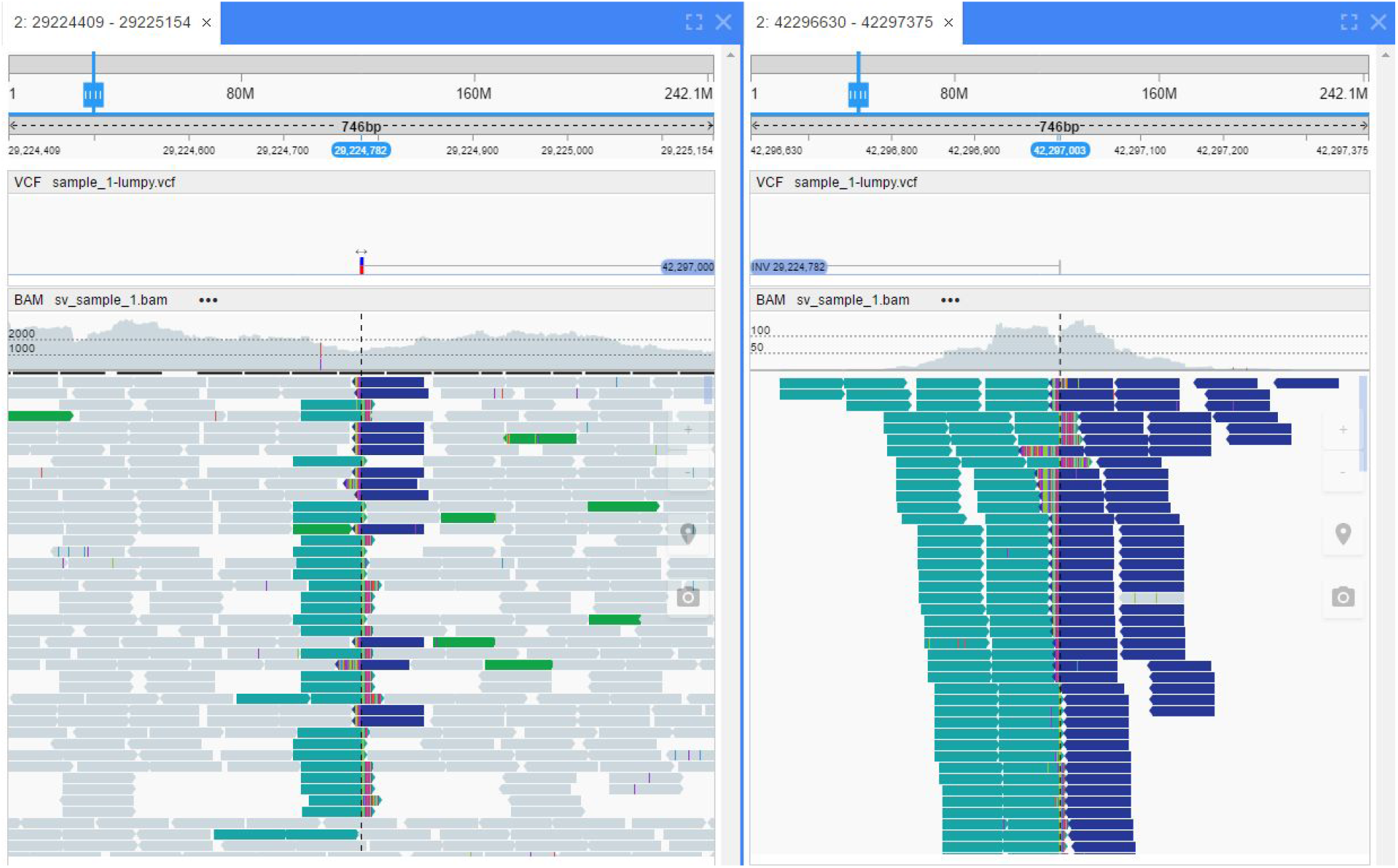
EML4-ALK inversion fusion. a. The effect of the fusion. b. The read evidence for the event at both breakpoints.

Other programs in development to better visualise the read level data include for example Genome Ribbon, see http://genomeribbon.com/.

## Conclusion and discussion

Here we presented a scheme for structural variant calling algorithms to prioritise for known fusion events as well as aberrations in a panel of cancer related genes.

This method prioritises based on biological information such as genes of interest and can be used in combination with orthogonal discovery based approaches [Ganel]. In their approach, Ganel et al. produce in silico SV impact predictions that can be useful when whittling down the number of SVs of unknown significance and narrowing down to the likely most pathogenic ones; if the more hypothesis driven prioritisation described here does not yield satisfactory results, the approach by [Ganel] might uncover additional novel variants.

We developed a visualisation framework in the New Genome Browser to illustrate the effects of the structural variants on genes in a user friendly, simple manner. We expect these visualisations to be extremely helpful for scientists in quickly producing publication ready gene fusion figures.

We look forward to accepting improvements from other groups to further improve structural variant calling interpretation and visualisation in cancer. This could be in the form of providing lists of genes of interest, suggesting alternative tiers to the prioritisation or adding support for other structural variant callers.

## Availability of data and algorithms

The vcf level data will be made available for all the samples except for the PDX model at https://github.com/AstraZeneca-NGS/. An implementation of the proposed methodology is available via bcbio, https://github.com/chapmanb/bcbio-nextgen. SnpEff 4.3 and later are available via http://snpeff.sourceforge.net/. The prioritisation code for structural variants is accessible at https://github.com/AstraZeneca-NGS/simple_sv_annotation. The New Genome Browser is available at https://github.com/epam/NGB. All software used herein is freely available under open source licences.

## Acknowledgements

The following centres are kindly acknowledged for sequencing the three replicates of the samples HDC134P, HDC140P and HDC141P.

West Midlands Regional Genetics Laboratory Birmingham Women’s NHS Foundation Trust Mindelsohn Way, Edgbaston Birmingham B15 2TG www.bwhct.nhs.uk/genetics-index/genetics-reglab-home.htm
Molecular Diagnostics The Centre for Molecular Pathology The Royal Marsden 15 Cotswold Road, Sutton Surrey SM2 5NG www.icr.ac.uk
All Wales Medical Genetics Service University Hospital of Wales Heath Park Cardiff CF14 4XW

## References

Can Alkan, Bradley P. Coe and Evan E. Eichler. Genome structural variation discovery and genotyping. Nature Reviews Genetics 12, 363-376 (2011)

T. Hedley Carr, Robert McEwen, Brian Dougherty, Justin H. Johnson, Jonathan R. Dry, Zhongwu Lai, Zara Ghazoui, Naomi M. Laing, Darren R. Hodgson, Francisco Cruzalegui, Simon J. Hollingsworth & J. Carl Barrett. Defining actionable mutations for oncology therapeutic development. Nature Reviews Cancer 16, 319–329 (2016)

Chen X, Schulz-Trieglaff O, Shaw R, Barnes B, Schlesinger F, Källberg M, Cox AJ, Kruglyak S, Saunders CT. Manta: rapid detection of structural variants and indels for germline and cancer sequencing applications. Bioinformatics. 2016 Apr 15;32(8):1220-2.

Cingolani P, Platts A, Wang le L, Coon M, Nguyen T, Wang L, Land SJ, Lu X, Ruden DM. A program for annotating and predicting the effects of single nucleotide polymorphisms, SnpEff: SNPs in the genome of Drosophila melanogaster strain w1118; iso-2; iso-3. Fly, 2012 Apr-Jun;6(2):80-92. PMID: 22728672

Eilbeck, Karen, et al. “The Sequence Ontology: a tool for the unification of genome annotations.” Genome biology 6.5 (2005): 1.

Liron Ganel, Haley J Abel, Ira M Hall. SVScore: An Impact Prediction Tool For Structural Variation. http://dx.doi.org/10.1101/073833

A sequence-level map of chromosomal breakpoints in the MCF-7 breast cancer cell line yields insights into the evolution of a cancer genome

Natasha S. Latysheva and M. Madan Babu. Discovering and understanding oncogenic gene fusions through data intensive computational approaches. Nucleic Acids Res. 2016 Jun 2;44(10):4487-503.

Ryan M Layer, Colby Chiang, Aaron R Quinlan and Ira M Hall. LUMPY: a probabilistic framework for structural variant discovery. Genome Biology 2014 15:R84.

Alexander Lex, Nils Gehlenborg, Hendrik Strobelt, Romain Vuillemot, Hanspeter Pfister, UpSet: Visualization of Intersecting Sets, IEEE Transactions on Visualization and Computer Graphics (InfoVis ‘14), vol. 20, no. 12, pp. 1983–1992, 2014.

Li, Heng. “Aligning sequence reads, clone sequences and assembly contigs with BWA-MEM.” arXiv preprint arXiv:1303.3997 (2013).

Márton Münz, Elise Ruark, Anthony Renwick, Emma Ramsay, Matthew Clarke, Shazia Mahamdallie, Victoria Cloke, Sheila Seal, Ann Strydom, Gerton Lunter, Nazneen Rahman. CSN and CAVA: variant annotation tools for rapid, robust next-generation sequencing analysis in the clinical setting. Genome Medicine 7:76, doi:10.1186/s13073-015-0195-6 (2015).

Daniel Nicorici, Mihaela Satalan, Henrik Edgren, Sara Kangaspeska, Astrid Murumagi, Olli Kallioniemi, Sami Virtanen, Olavi Kilkku. FusionCatcher - a tool for finding somatic fusion genes in paired-end RNA-sequencing data. bioRxiv, Nov. 2014, DOI:10.1101/011650

Aaron C Noll, Neil A Miller, Laurie D Smith, Byunggil Yoo, Stephanie Fiedler, Linda D Cooley, Laurel K Willig, Josh E Petrikin, Julie Cakici, John Lesko, Angela Newton, Kali Detherage, Isabelle Thiffault, Carol J Saunders, Emily G Farrow & Stephen F Kingsmore. Clinical detection of deletion structural variants in whole-genome sequences [Veeraraghavan] Recurrent ES R1-CCDC170 rearrangements in an aggressive subset of estrogen-receptor positive breast cancers

Manabu Soda, Young Lim Choi, Munehiro Enomoto, Shuji Takada, Yoshihiro Yamashita, Shunpei Ishikawa, Shin-ichiro Fujiwara, Hideki Watanabe, Kentaro Kurashina, Hisashi Hatanaka, Masashi Bando, Shoji Ohno, Yuichi Ishikawa, Hiroyuki Aburatani, Toshiro Niki, Yasunori Sohara, Yukihiko Sugiyama & Hiroyuki Mano. Identification of the transforming EML4–ALK fusion gene in non-small-cell lung cancer. Nature 448, 561-566 (2007).

Spies N, Zook JM, Salit M, Sidow A. 2015. svviz: a read viewer for validating structural variants. Bioinformatics doi:bioinformatics/btv478

N Sugawa, A J Ekstrand, C D James, and V P Collins. Identical splicing of aberrant epidermal growth factor receptor transcripts from amplified rearranged genes in human glioblastomas. Proc Natl Acad Sci U S A. 1990 Nov; 87(21): 8602–8606.

Lorenzo Tattini, Romina D’Aurizio, and Alberto Magi. Detection of Genomic Structural Variants from Next-Generation Sequencing Data. Front Bioeng Biotechnol. 2015; 3: 92.

Scott A Tomlins, Bharathi Laxman, Sooryanarayana Varambally, Xuhong Cao, Jindan Yu, Beth E Helgeson, Qi Cao, John R Prensner, Mark A Rubin, Rajal B Shah, Rohit Mehra, and Arul M Chinnaiyan. Role of the TMPRSS2-ERG Gene Fusion in Prostate Cancer. Neoplasia. 2008 Feb; 10(2): 177–188.

The Cancer Genome Atlas Research Network. Comprehensive molecular characterization of urothelial bladder carcinoma. Nature 507, 315–322

